# Linkage mapping of yeast cross protection connects gene expression variation to a higher-order organismal trait

**DOI:** 10.1101/152793

**Authors:** Tara N. Stuecker, Amanda N. Scholes, Jeffrey A. Lewis

## Abstract

Gene expression variation is extensive in nature, and is hypothesized to play a major role in shaping phenotypic diversity. However, connecting differences in gene expression across individuals to higher-order organismal traits is not trivial. In many cases, gene expression variation may be evolutionarily neutral, and in other cases expression variation may only affect phenotype under specific conditions. To understand connections between gene expression variation and stress defense phenotypes, we have been leveraging extensive natural variation in the gene expression response to acute ethanol in laboratory and wild *Saccharomyces cerevisiae* strains. Previous work found that the genetic architecture underlying these expression differences included dozens of “hotspot” loci that affected many transcripts in *trans*. In the present study, we provide new evidence that one of these expression QTL hotspot loci is responsible for natural variation in one particular stress defense phenotype—ethanol-induced cross protection against severe doses of H_2_O_2_. The causative polymorphism is in the heme-activated transcription factor Hap1p, which we show directly impacts cross protection, but not the basal H_2_O_2_ resistance of unstressed cells. This provides further support that distinct cellular mechanisms underlie basal and acquired stress resistance. We also show that the Hap1p-dependent cross protection relies on novel regulation of cytosolic catalase T (Ctt1p) during ethanol stress in wild strains. Because ethanol accumulation precedes aerobic respiration and accompanying reactive oxygen species formation, wild strains with the ability to anticipate impending oxidative stress would likely be at an advantage. This study highlights how strategically chosen traits that better correlate with gene expression changes can improve our power to identify novel connections between gene expression variation and higher-order organismal phenotypes.

## Introduction

A fundamental question in genetics is how individuals with extremely similar genetic makeups can have dramatically different characteristics. One hypothesis is that a small number of regulatory polymorphisms can have large effects on gene expression, leading to the extensive phenotypic variation we see across individuals. In fact, gene expression variation is hypothesized to underlie the extensive phenotypic differences we see between humans and chimpanzees despite >98% DNA sequence identity [1, 2]. This hypothesis is supported by numerous examples of gene expression variation affecting higher-order organismal traits. For example, human genome-wide association studies (GWAS) have found that a substantial fraction of disease-associated variants are concentrated in non-coding regulatory DNA regions [3–8]. Further examples include gene expression variation being linked to differences in metabolism [9–11], physiology [12–16], morphology [17–23], and behavior [24–27].

While gene expression variation is pervasive, there is often a lack of obvious phenotypic change associated with differentially expressed genes. This can occur for a variety of reasons. First, a large fraction of expression variation has been postulated to be evolutionarily neutral with no effect on organismal fitness [28–30]. Second, co-regulation of genes that share the same upstream signaling network and transcription factors can lead to genes whose expression differences correlate with phenotype but are not truly causative. Finally, some gene expression differences may truly affect phenotype, but only under specific conditions. For example, the predictive power of expression quantitative trait loci (eQTL) mapping studies on higher-order phenotypes can be poor unless multiple environments are considered [31]. Similarly, tissue-restricted eQTLs are more likely to map to known disease-associated loci identified from GWAS than non-tissue-restricted eQTLs [32, 33].

Thus, a major challenge for connecting gene expression variation to downstream effects on higher-order traits is the choice of which conditions and traits to examine. To this end, we have been leveraging natural variation in the model eukaryote *Saccharomyces cerevisiae*, and a phenotype called acquired stress resistance. Many studies have shown a poor correlation between genes that respond to stress and their importance for surviving stress [34–43]. Thus, we and others have argued that the role of stress-activated gene expression is not to survive the initial insult, but instead protects cells from impending severe stress through a phenomenon called acquired stress resistance [44, 45]. Acquired stress resistance (sometimes referred to as “induced tolerance” or the “adaptive response”) occurs when cells pretreated with a mild dose of stress gain the ability to survive an otherwise lethal dose of severe stress. Notably, acquired stress resistance can occur when the mild and severe stresses are the same (same-stress protection) or across pairs of different stresses (cross protection). This phenomenon has been observed in diverse organisms ranging from bacteria to higher eukaryotes including humans [44–50]. The specific mechanisms governing acquisition of higher stress resistance are poorly understood, but there are wide reaching implications. In humans, ischemic preconditioning (transient ischemia followed by reperfusion—i.e. mild stress pretreatment followed by severe stress) may improve outcomes of cardiovascular surgery [51–54], while transient ischemic attacks (“mini-strokes”) may protect the brain during massive ischemic stroke [55–57]. Thus, understanding the genetic basis of acquired stress resistance in model organisms holds promise for mitigating the effects of stress in humans.

A previous study found that a commonly used S288c lab strain is unable to acquire further ethanol resistance when pretreated with a mild dose of ethanol [44]. We found this phenotype to be surprising, considering the unique role ethanol plays in the life history of *Saccharomyces* yeast, where the evolution of aerobic fermentation gave yeast an advantage over ethanol-sensitive competitors [58]. Because ethanol is a self-imposed stress that induces a robust stress response [59–63], we expected that ethanol should provoke acquired stress resistance in wild yeast strains. Indeed, this turned out to be the case, with the majority of tested wild strains acquiring resistance to severe ethanol following a mild ethanol treatment [45]. Furthermore, this phenotype correlated with extensive differences in the transcriptional response to acute ethanol stress in the lab strain when compared to a wild vineyard (M22) and wild oak (YPS163) strain (>28% of S288c genes were differentially expressed at an FDR of 0.01) [45, 64]. We performed linkage mapping of S288c crossed to a wild vineyard strain (M22) and wild oak strain (YPS163), and observed numerous “hotspots” where the same eQTL loci affects the expression of a large number of transcripts (anywhere from 10 – 500 transcripts per hotspot) [64].

In the present study, we provide new evidence that one of these eQTL hotspot loci is responsible for natural variation in acquired stress resistance, namely the ability of ethanol to cross protect against oxidative stress in the form of hydrogen peroxide. The causative polymorphism is in the heme-activated transcription factor Hap1p, which we show directly impacts cross protection, but not the basal resistance of unstressed cells. Finally, we show that the Hap1p effect is mediated through novel regulation of cytosolic catalase T (Ctt1p) during ethanol stress in wild strains. This study highlights how strategically chosen traits that best correlate with gene expression changes can improve our power to identify novel connections between gene expression variation and higher-order organismal phenotypes.

## Results

### The Genetic Basis of Natural Variation in Yeast Cross Protection

We previously found that an S288c-derived lab strain was unable to acquire further ethanol resistance when pretreated with a mild dose of ethanol, in contrast to the vast majority of ~50 diverse yeast strains [45]. In addition to the S288c strain’s acquired ethanol resistance defect, ethanol also failed to cross protect against other subsequent stresses [44, 65]. In nature, wild yeast cells ferment sugars to ethanol, and then shift to a respiratory metabolism that generates endogenous reactive oxygen species [66–68]. Thus, we hypothesized that ethanol might cross protect against oxidative stress in wild yeast strains. We tested this hypothesis by assessing whether mild ethanol treatment would protect a wild oak strain (YPS163) from severe oxidative stress in the form of hydrogen peroxide (H_2_O_2_). Cross protection assays were performed by exposing cells to a mild, sublethal dose of ethanol (5% v/v) for 60 min, followed by exposure to a panel of 11 increasingly severe doses of H_2_O_2_ (see Materials and Methods). Confirming the observations of Berry and Gasch [44], ethanol failed to cross protect against H_2_O_2_ in S288c, and in fact slightly exacerbated H_2_O_2_ toxicity (Fig 1). In contrast, ethanol strongly cross protected against H_2_O_2_ in YPS163 (Fig 1).

**Figure 1.**
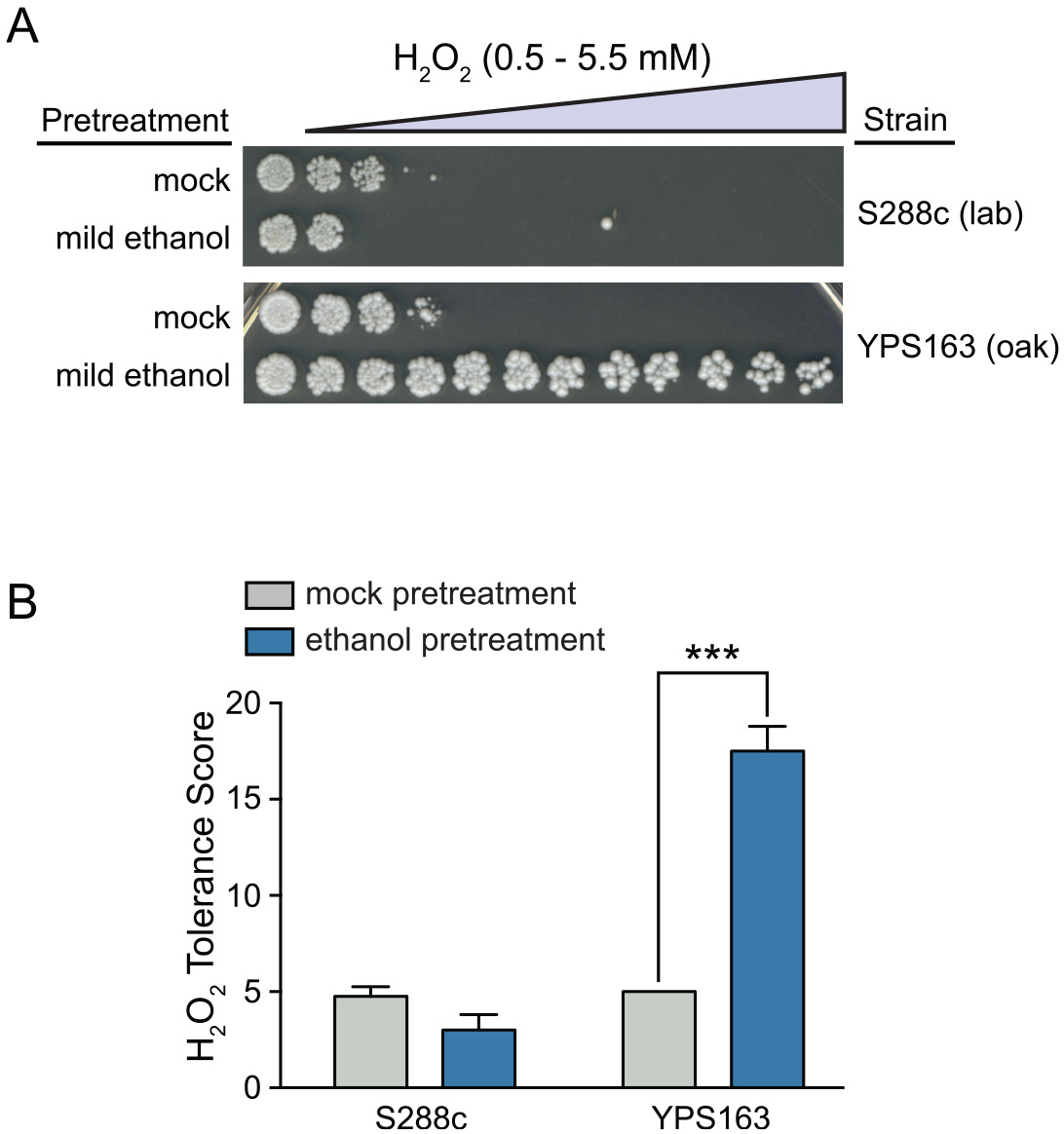
Natural variation in ethanol-induced cross protection against H_2_O_2_. (A) A representative acquired H_2_O_2_ resistance assay is shown. S288c (lab strain) and YPS163 (wild oak strain) were exposed to 5% ethanol or mock (5% water) pretreatment for 60 min, washed, exposed to 11 doses of severe H_2_O_2_ for 2 hr, and then plated to score viability. (B) A single survival score was calculated from the viability at all H_2_O_2_ doses (see Materials and Methods). Each plot shows the mean and standard deviation of 4 independent biological replicates. Asterisks represent resistance that was significantly different from mock-treated cells (*** *P* < 0.001, *t*-test).

The inability of ethanol to induce acquired stress resistance in S288c correlates with thousands of differences in ethanol-dependent gene expression in comparison to wild strains that can acquire ethanol resistance [45, 64]. In light of this observation, and the known dependency of cross protection on stress-activated gene expression changes [44], we hypothesized that differences in cross protection against H_2_O_2_ by ethanol may be linked to differential gene expression. To test this, we performed quantitative trait loci (QTL) mapping using the same mapping population as our original eQTL study that mapped the genetic architecture of ethanol-responsive gene expression [64]. Specifically, we conducted QTL mapping of both basal and acquired H_2_O_2_ resistance in 44 F_2_ progeny of S288c crossed with YPS163 (see Materials and Methods). While we found no significant QTLs for basal H_2_O_2_ resistance, we did find a significant QTL peak on chromosome XII for cross protection (Fig 2). It is unlikely that our failure to detect a chromosome XII QTL for basal H_2_O_2_ resistance was due to a lack of statistical power, because two independent basal H_2_O_2_ resistance QTL studies using millions of S288c x YPS163 F_2_ segregants also found no significant associations at this locus [69, 70]. Additionally, we estimated the heritability of phenotypic variation in basal resistance to be 0.79, which is slightly above the median value estimated by Bloom and colleagues for 46 yeast traits [71], and is only moderately lower than the heritability for cross protection (0.92). Thus, it is likely that the genetic basis of natural variation in acquired stress resistance is distinct from the basal resistance of unstressed cells (see Discussion).

**Figure 2.**
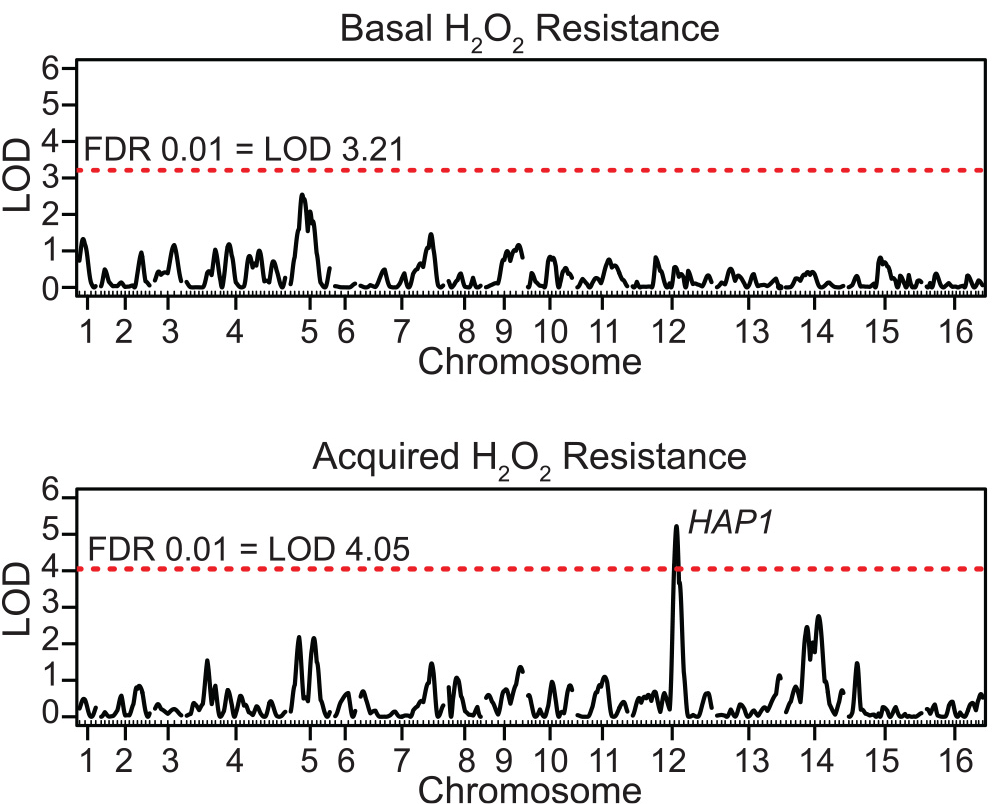
The genetic basis of natural variation for basal and acquired stress resistance is distinct. Linkage mapping of the S288c = YPS163 cross identified no significant QTLs for basal H_2_ O_2_ resistance (top panel), but did identify a major QTL on chromosome XII for ethanol-induced cross protection against H_2_O_2_ (bottom panel). *HAP1* falls within the 1.5-LOD support interval for the chromosome XII peak. The red horizontal line denotes the LOD threshold for significance (1% FDR).

The significant QTL for cross protection was located near a known polymorphism in *HAP1*, a heme-dependent transcription factor that controls genes involved in aerobic respiration [72–74], sterol biosynthesis [75–77], and interestingly, oxidative stress [77, 78]. S288c harbors a known defect in *HAP1*, where a Ty1 transposon insertion in the 3’ end of the gene’s coding region has been shown to reduce its function [79]. In fact, we previously hypothesized that the defective *HAP1* allele was responsible for the inability of S288c to acquire further resistance to ethanol. However, a YPS163 *hap1Δ* strain was still fully able to acquire ethanol resistance, despite notable differences in the gene expression response to ethanol in the mutant [45]. Likewise, despite previous studies implicating Hap1p as a regulator of oxidative stress defense genes [77, 78], *HAP1* is apparently dispensable for same-stress acquired H_2_O_2_ resistance [47]. These observations suggest that the molecular mechanisms underlying various acquired stress resistance phenotypes can differ, even when the identity of the secondary stress is the same.

### Ethanol fails to cross protect against severe H_2_O_2_ in strains lacking *HAP1* function

Because we previously implicated *HAP1* as a major ethanol-responsive eQTL hotspot affecting over 100 genes, we hypothesized that ethanol-induced cross protection against H_2_O _2_ may depend upon Hap1p-regulated genes. However, it was formally possible that *HAP1* was merely linked to the truly causal polymorphism. Thus, we performed a series of experiments to definitively test whether the polymorphism in *HAP1* was causative for the S288c strain’s inability to acquire H_2_O_2_ resistance following ethanol pretreatment. First, we deleted *HAP1* in the YPS163 background and found that the YPS163 *hap1*Δ mutant had highly diminished acquired H_2_O_2_ resistance (Figs 3B and 3C). Notably, the diminished acquired H_2_O_2_ resistance of the YPS163 *hap1*Δ mutant was still higher than that of the S288c strain, suggesting that the Ty1 insertion into S288c *hap1* allele may produce more severe effects than a deletion, and may in fact operate as dominant-negative mutation. However, a YPS163-S288c hybrid containing both *HAP1* alleles fully acquired H_2_O_2_ resistance (Figs 3B and 3C), suggesting that *HAP1^S288c^* is recessive.

**Figure 3.**
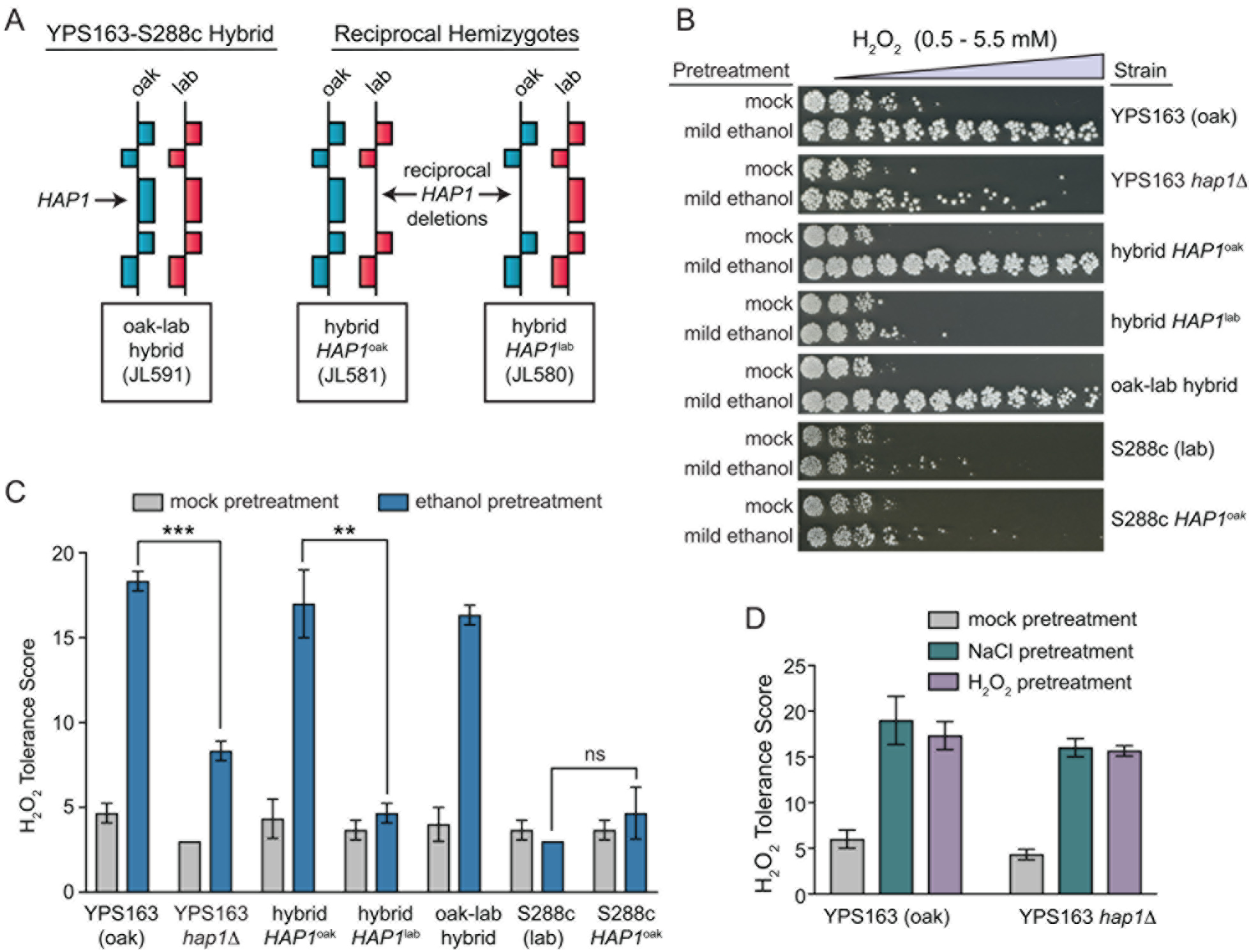
Ethanol-induced cross protection against H_2_O_2_ requires *HAP1.* (A) Schematic of reciprocal hemizygosity analysis. Each block represents a gene, and each hybrid strain contains a single-copy deletion of *hap1*, and a single copy of either the *HAP1^S288c^* (lab) or *HAP1^YPS163^* (oak) allele. (B) Representative acquired H_2_O_2_ resistance assays for wild-type YPS163 (oak), YPS163 *hap1*Δ mutant, reciprocal hemizygotes, S288c (lab), and S288c repaired with the *HAP1^YPS163^* allele. (C) Each survival score plot shows the mean and standard deviation of biological triplicates. Asterisks represent significant differences in acquired resistance between denoted strains (** *P* < 0.01, *** *P* < 0.001, ns = not significant (*P* > 0.05), *t*-test). (D) *HAP1* is not required for acquired H_2_O_2_ resistance following mild H_2_O_2_ or mild NaCl pretreatments. YPS163 and the YPS163 *hap1*Δ mutant scored for acquired H_2_O_2_ resistance following mild pretreatment with either 0.4 mM H_2_O_2_ or 0.4 M NaCl. The survival score plot shows the mean and standard deviation of biological triplicates.

Second, we applied an approach called reciprocal hemizygosity analysis [80], where each *HAP1* allele is analyzed in an otherwise isogenic S288c-YPS163 hybrid background (see Fig 3A for a schematic). In each of the two reciprocal strains, one allele of *HAP1* is deleted,
producing a hybrid strain containing either the S288c or YPS163 *HAP1* allele in single copy (i.e. hemizygous for *HAP1*). We found that the hybrid strain containing the *HAP1 ^YPS163^* allele showed full cross protection, while the strain containing the *HAP1^S288c^* allele showed none (Fig 3B and 3C). There was no difference in basal stress resistance in the reciprocal strains, providing an additional line of evidence that the genetic basis of basal and acquired stress resistance is distinct. Furthermore, the YPS163 *hap1*Δ mutant was unaffected for acquired H_2_O_2_ resistance when mild H_2_O_2_ or mild NaCl were used as mild stress pretreatments (Fig 3D), suggesting that Hap1p plays a distinct role in ethanol-induced cross protection (see Discussion).

Finally, we tested whether repair of the defective *hap1* allele in S288c could restore cross protection. Surprisingly, S288c repaired with *HAP1 ^YPS163^* was largely unable to acquire further H_2_O _2_ resistance (Figs 3B and 3C). This additional layer of genetic complexity suggests that S288c harbors additional polymorphisms that affect cross protection. Moreover, these alleles are apparently masked in YPS163-S288c hybrids that fully acquire H_2_O_2_ resistance, suggesting that they are recessive (see Discussion).

### Strains lacking *HAP1* function show decreased catalase expression and reduced peroxidase activity during ethanol stress

Because Hap1p is a transcription factor, we hypothesized that acquired H_2_O _2_ resistance relied on Hap1p-dependent expression of a stress protectant protein. We reasoned that the putative stress protectant protein should have the following properties: i) a biological function consistent with H_2_O_2_ detoxification or damage repair, ii) reduced ethanol-responsive expression in S288c versus YPS163, iii) be a target gene of the *HAP1* eQTL hotspot, and iv) possess evidence of regulation by Hap1p.

We first looked for overlap between our previously identified *HAP1* eQTL hotspot (encompassing 376 genes) and genes with significantly reduced ethanol-responsive induction in S288c versus YPS163 (309 genes) [64]. Thirty-four genes overlapped for both criteria, including several that directly defend against reactive oxygen species (*TSA2* encoding thioredoxin peroxidase, *SOD2* encoding mitochondrial manganese superoxide dismutase, *CTT1* encoding cytosolic catalase T, and *GSH1* encoding *γ*-glutamylcysteine synthetase (Fig 4A and S1 Table)). Of those 34 genes, 8 also had direct evidence of Hap1p binding to their promoters [81] (Fig 4B and S1 Table), including *CTT1* and *GSH1* (though both *TSA2* and *SOD2* have indirect evidence of regulation by Hap1p [82, 83]).

**Figure 4.**
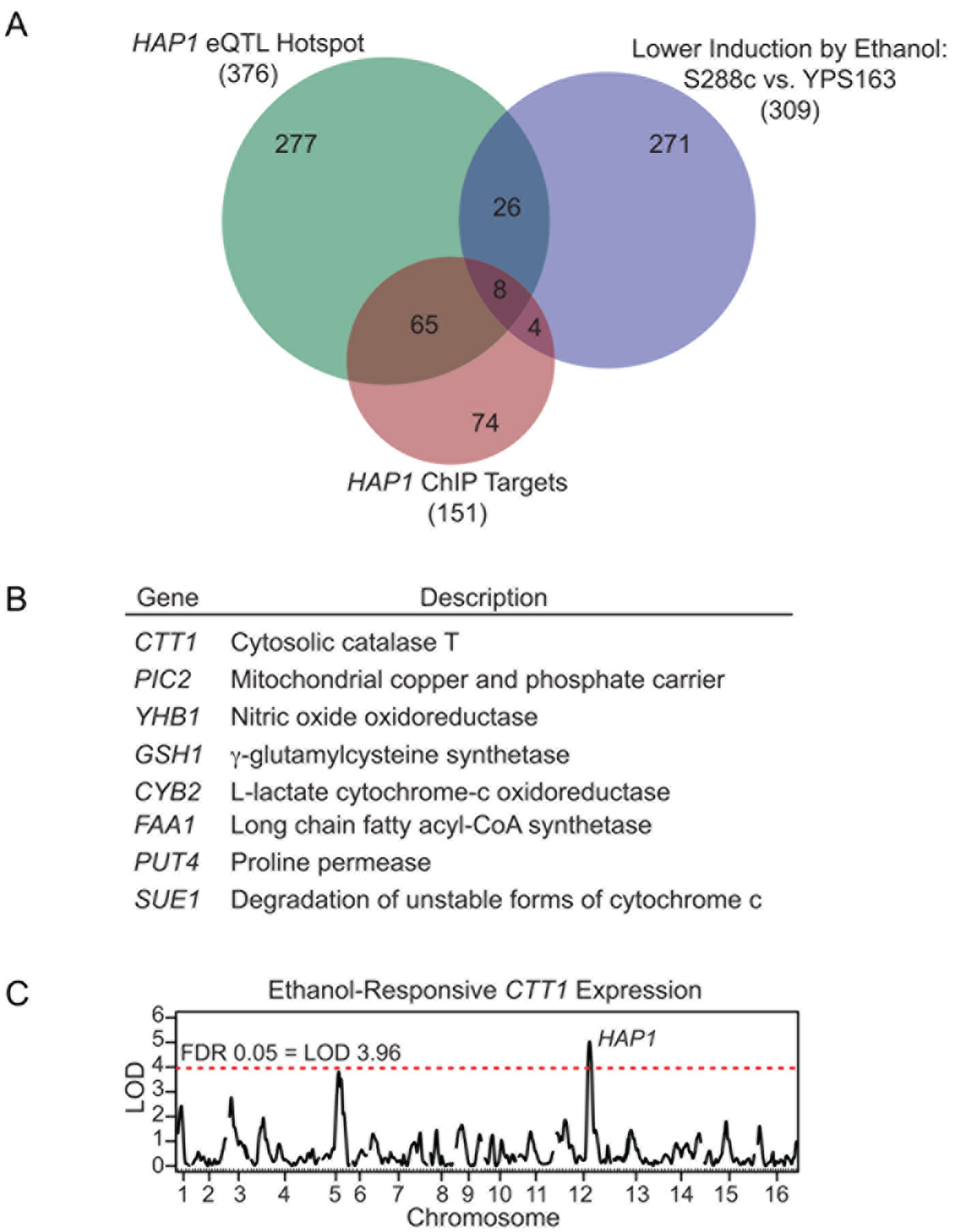
Expression variation in Hap1p regulatory targets implicates oxidative stress defense genes as the direct effectors of ethanol-induced cross protection against H_2_O_2_. (A) Overlap between genes that were *HAP1* eQTL hotspot targets from [64], genes with defective induction in S288c vs. YPS163 from [64], and direct targets of *HAP1* identified via ChIP experiments compiled from [81]. (B) Descriptions of the eight genes that overlapped for all three criteria. (C) Previous eQTL mapping of the yeast ethanol response (newly plotted here using data described in [64]), implicated *HAP1* as causative for natural variation in *CTT1* induction levels during ethanol stress.

We first focused on *CTT1*, since it is both necessary for NaCl-induced cross protection against H_2_O_2_ in S288c [84], and sufficient to increase H_2_O_2_ resistance when exogenously overexpressed in S288c [85]. We deleted *CTT1* in the YPS163 background, and found that ethanol-induced cross protection against H _2_O_2_ was completely eliminated (Fig 5). The complete lack of cross protection in the *ctt1*Δ mutant suggests that other peroxidases cannot compensate for the lack of catalase activity under this condition. Next, because *CTT1* was part of the *HAP1* eQTL hotspot (Fig 4C, plotted using the data described in [64]), we tested whether the *HAP1^S288c^* allele reduced *CTT1* expression during ethanol stress. To do this, we performed qPCR to measure *CTT1* mRNA induction following a 30-minute ethanol treatment (i.e. the peak ethanol response [45]). Consistent with our previous microarray data [45, 64], we saw lower induction of *CTT1* by ethanol in S288c relative to YPS163 (Fig 6A). Moreover, we saw dramatically reduced induction of *CTT1* in a YPS163 *hap1*Δ mutant compared to the wild-type YPS163 control (Fig 6A). Further support that *HAP1* is causative for reduced *CTT1* expression was provided by performing qPCR in the *HAP1* reciprocal hemizygotes, where we found that the *HAP1^S288c^* allele resulted in significantly reduced *CTT1* induction compared to the *HAP1^YPS163^* allele (Fig 6A).

**Figure 5.**
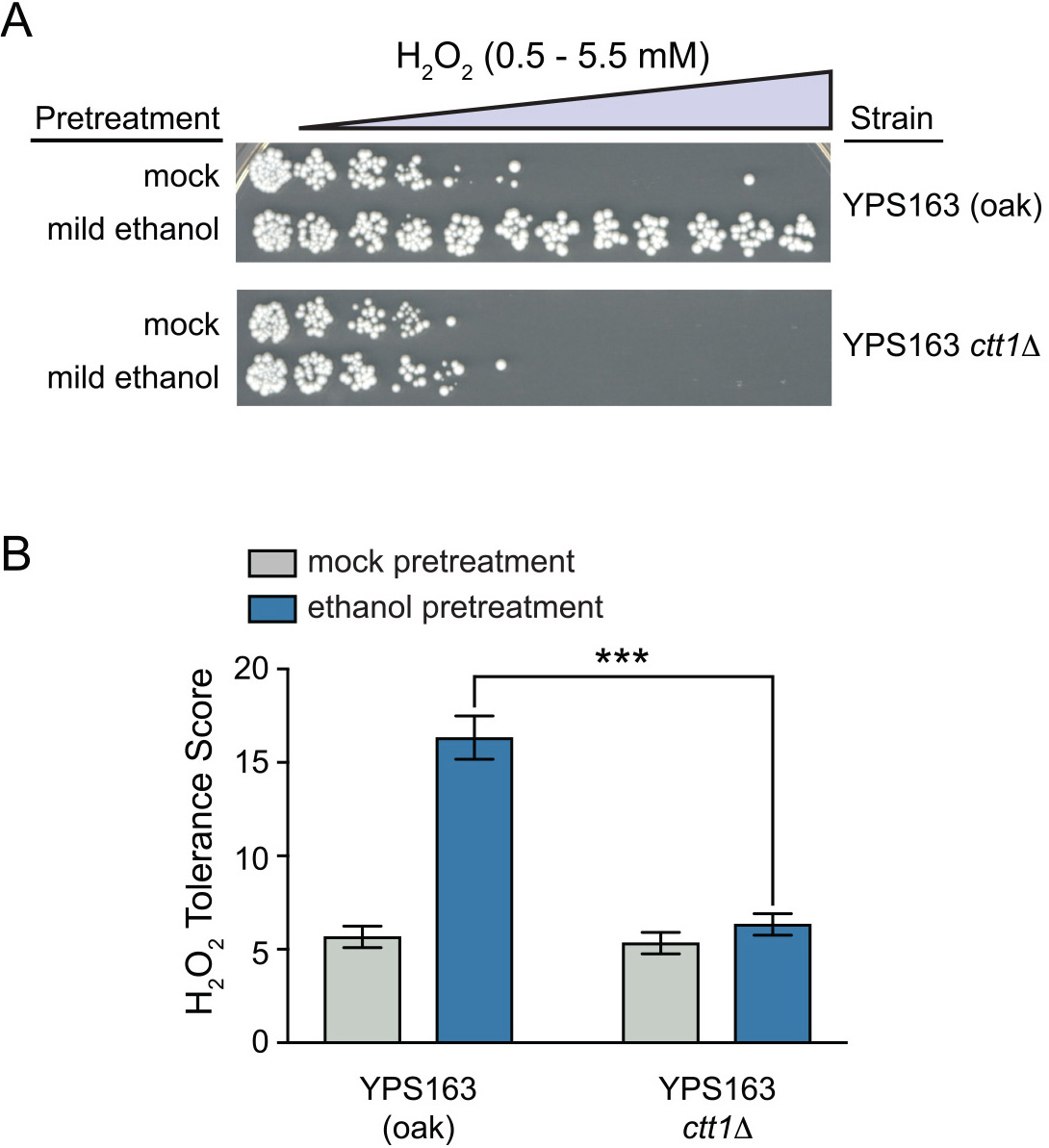
*CTT1* function is necessary for ethanol-induced cross protection against H_2_O_2_. (A) Representative acquired H_2_O_2_ resistance assays for wild-type YPS163 and the YPS163 *ctt1*Δ mutant. (B) Survival score plots indicating the mean and standard deviation of biological triplicates. Asterisks represent significant differences in acquired resistance between denoted strains (*** *P* < 0.001, *t*-test).

To determine whether the differences in *CTT1* induction across strain backgrounds also manifested as differences in each strain’s ability to detoxify H_2_O_2_, we measured *in vitro* peroxidase activity in cell-free extracts. We compared *in vitro* peroxidase activity in extracts from unstressed cells and cells exposed to ethanol stress for 60 minutes (i.e. the same pre-treatment time that induces acquired H_2_O_2_ resistance (see Materials and Methods)). For wild-type YPS163, ethanol strongly induced peroxidase activity, and this induction was completely dependent upon *CTT1* (Fig 6b). Mirroring *CTT1* gene expression patterns, the induction of peroxidase activity was reduced in a YPS163 *hap1*Δ mutant. Additionally, reciprocal hemizygosity analysis provided further support that lack of *HAP1* function results in decreased peroxidase activity, as the hybrid containing the *HAP1^S288c^* allele showed significantly reduced peroxidase activity following ethanol stress compared to the hybrid containing the *HAP1^YPS163^* allele (Fig 6b). Notably, the hybrid containing the *HAP1^YPS163^* allele had lower *CTT1* induction and *in vitro* peroxidase activity following ethanol shock than wild-type YPS163, despite equivalent levels of acquired H_2_O_2_ resistance in the strains. These results suggest that *HAP1* may play additional roles in acquired H_2_O_2_ resistance beyond H_2_O_2_ detoxification, depending upon the genetic background (see Discussion). Interestingly, S288c showed no induction of peroxidase activity upon ethanol treatment, despite modest induction of the *CTT1* transcript. This result is reminiscent of Ctt1p regulation during heat shock in the S288c background, where mRNA levels increase without a concomitant increase in protein levels [84]. Thus, in addition to strain-specific differences in *CTT1* regulation at the RNA level, there are likely differences in regulation at the level of translation and/or protein stability.

**Figure 6.**
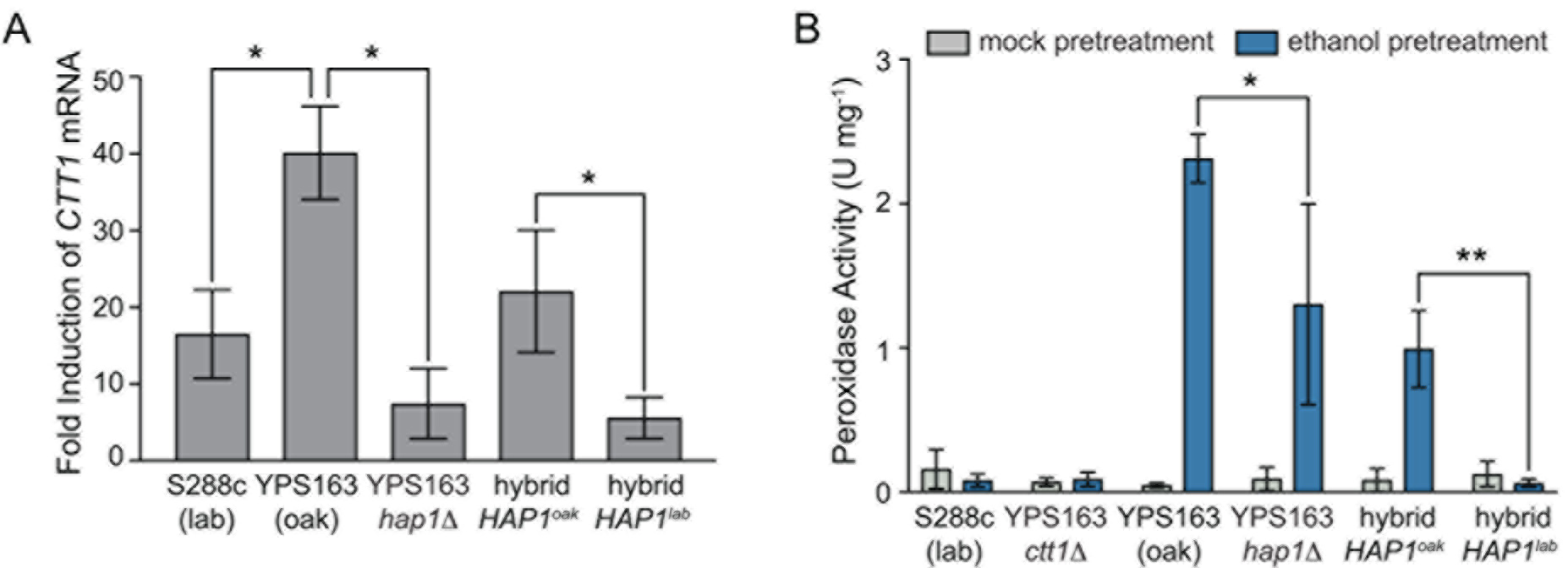
*HAP1* is required for full induction of *CTT1* gene expression and cellular peroxidase activity during ethanol stress. (A) Fold induction of *CTT1* mRNA in indicated strains following 30 min ethanol stress compared to unstressed cells, assessed by qPCR. (B) Peroxidase activity measured in cell-free extracts in either mock-treated or ethanol-stressed cells. The plots indicate the mean and standard deviation of biological triplicates (mRNA) or quadruplicates (peroxidase activity). Asterisks represent significant differences in *CTT1* mRNA induction or peroxidase activity between denoted strains (* *P* < 0.05, ** *P* < 0.01, paired *t*-test).

## Discussion

In this study, we leveraged extensive natural variation in the yeast ethanol response to understand potential connections between gene expression variation and higher-order organismal traits. Previous screens of gene deletion libraries have found surprisingly little overlap between the genes necessary for surviving stress and genes that are induced by stress. [34–43]. Instead, gene induction may be a better predictor of a gene’s requirement for acquired stress resistance [84]. Thus, we hypothesized that phenotypic variation in acquired stress resistance may be linked to natural variation in stress-activated gene expression. Our results provide a compelling case study in support of this notion—namely that a polymorphism in the *HAP1* transcription factor is causative for variation in acquired H_2_O_2_ resistance, but not for the basal H_2_O_2_ resistance of unstressed cells. Forward genetic screens have shown that the genes necessary for basal and acquired resistance are largely non-overlapping [34, 36, 84], suggesting that mechanisms underlying basal and acquired stress resistance are distinct. That the YPS163 *hap1*Δ mutant was only affected for acquired H_2_O_2_ resistance, but not the basal resistance of unstressed cells, strongly supports this model. Moreover, the YPS163 *hap1*Δ mutant was affected only when ethanol was the mild pretreatment, and was able to fully acquire H_2_O_2_ resistance following mild H_2_O_2_ or mild NaCl (Fig 3D). These results suggest that the mechanisms underlying acquired resistance differ depending upon the mild stress that provokes the response. Further dissection of the mechanisms underlying acquired stress resistance will provide a more integrated view of eukaryotic stress biology.

Our results reveal a new role for Hap1p in cross protection against H _2_O_2_ that has been lost in the S288c lab strain. We propose that a major mechanism underlying ethanol-induced cross protection against H_2_O_2_ is the induction of cytosolic catalase T (Ctt1p), and that Hap1p is necessary for proper induction of *CTT1* during ethanol stress. We based this mechanism on the following observations. First, over-expression of *CTT1* in S288c is sufficient to induce high H _2_O_2_ resistance [85]. Second, a YPS163 *ctt1*Δ mutant cannot acquire any further H_2_O_2_ resistance following ethanol pre-treatment (Fig. 5), suggesting that no other antioxidant defenses are able to compensate under this condition. Lastly, the defect in cross protection for the YPS163 *hap1*Δ mutant correlates with reduced *CTT1* expression and peroxidase activity during ethanol stress (compare Figs 3 and 6). How Hap1p is involved in the regulation of *CTT1* during ethanol stress remains an open question, but we offer some possibilities. Hap1p is activated by heme, thus promoting transcription of genes involved in respiration, ergosterol biosynthesis, and oxidative stress defense including *CTT1* [75, 76, 78, 82]. Because heme biosynthesis requires oxygen, Hap1p is an indirect oxygen sensor and regulator of aerobically expressed genes [74, 75, 86]. There is currently no evidence that heme levels are affected by ethanol stress, nor is there evidence that Hap1p is “super-activating” under certain conditions. Thus, we disfavor a mechanism of induction caused solely by Hap1p activation. Instead, we favor a mechanism where Hap1p interacts with other transcription factors at the *CTT1* promoter during ethanol stress, leading to full *CTT1* induction. One possibility that we favor is recruitment of the general stress transcription factor Msn2p, which plays a known role in acquired stress resistance [44, 45]. We previously showed that a YPS163 *msn2*Δ mutant had no induction of *CTT1* mRNA during ethanol stress [45], suggesting that Msn2p was an essential activator for *CTT1* under this condition. The *CTT1* promoter region contains three Msn2p DNA-binding sites, two of which are ~100-bp away from the Hap1p binding site. Hap1p binding to the *CTT1* promoter could help recruit Msn2p during ethanol stress, possibly through chromatin remodeling that increases accessibility of the Msn2 binding sites as proposed by Elfving and colleagues [87].

What is the physiological role of Hap1p- dependent induction of *CTT1* during ethanol stress? One possibility is that regulation tied to the heme- and oxygen-sensing role of Hap1p ensures that *CTT1* induction only occurs under environmental conditions where reactive oxygen species (ROS) are most likely to be encountered—namely stressful conditions that are also aerobic. In the context of ethanol stress, aerobic fermentation would lead to subsequent respiration of the produced ethanol and simultaneous ROS production. Under these conditions, *CTT1* induction leading to ethanol-mediated cross protection against ROS would likely confer a fitness advantage. On the other hand, during stressful yet anoxic conditions, Ctt1p and other ROS-scavenging proteins are likely unnecessary. Furthermore, because heme is not synthesized during anoxic conditions [74], Hap1p fails to induce *CTT1* and other genes encoding non-essential heme-containing proteins. This may improve fitness by conserving energy used for biosynthesis and by redirecting limited heme to more essential heme-containing proteins.

The S288c lab strain has long been known to possess a defective *HAP1* allele [79]. Apparently, the defective allele arose relatively recently, as only S288c contains a *HAP1* Ty1 insertion out of over 100 sequenced strains [88, 89]. The lack of *HAP1* function in S288c could be due to relaxation of selective constraint, though others have argued in favor of positive selection for reduced ergosterol biosynthetic gene expression [90, 91]. Regardless, the loss of ethanol-induced acquired H_2_O_2_ resistance is likely a secondary effect of the loss of Hap1p function. Intriguingly, we did find that two (non- S88c) domesticated yeast strains also lack ethanol-induced cross protection against H_2_O_2_ (S1 Fig), suggesting that phenotypic differences in acquired stress resistance may differentiate domesticated versus wild yeast. Because environmental stresses are likely encountered in combination or sequentially [92], acquired stress resistance is likely an important phenotype in certain natural ecological settings. Future studies directed at understanding differences in acquired stress resistance phenotypes in diverse wild yeast strains may provide unique insights into the ecology of yeast.

While our QTL mapping identified *HAP1* as the major effector of cross protection, we note that additional complexity remains unexplained. Notably, despite the strong cross protection defect in the YPS163 *hap1*Δ mutant, some residual cross protection persists that is absent in S288c (compare Figs 1 and 3). Intriguingly, the residual cross protection is also absent in the hybrid carrying the *HAP1^S288c^* allele, suggesting the involvement of other genes depending upon the genetic background (Figs 3B and 3C). The lack of cross protection in S288c and the *HAP1^S288c^* hybrid correlates with the lack of inducible peroxidase activity following ethanol pretreatment in those strains. The lack of inducible peroxidase activity in S288c despite modest induction of *CTT1* mRNA could be due to translational regulation, as suggested by the observation that while mild heat shock induces *CTT1* mRNA, protein levels remain nearly undetectable [84]. Additionally, the hybrid carrying the *HAP1^YPS163^* allele still cross protects despite levels of *CTT1* mRNA induction and peroxidase activity that are lower than in YPS163, suggesting that other genes and processes are involved in this complex trait. This is strongly supported by the lack of complementation by the *HAP1^YPS163^* allele in the S288c background, which points to the presence of additional loci in S288c that affect acquired stress resistance. It is known that yeast strains with respiratory defects have increased ROS sensitivity [93, 94], potentially due to increased programmed cell death [95]. It is possible that reduced respiratory activity and concomitant ROS sensitivity in strains lacking *HAP1* is exacerbated by genetic interactions with other alleles. Future high resolution mapping experiments will be necessary to identify and characterize the source of these genetic background effects.

Gene expression variation is extensive in nature and is hypothesized to be a major driver of higher-order phenotypic variation. However, there are inherent challenges to connecting gene expression variation to higher-order organismal traits. Hundreds to thousands of genes are often differentially expressed across individuals, so identifying which particular transcripts exert effects on fitness is difficult. By studying acquired stress resistance—a phenotype better correlated with stress-activated gene expression changes—we were able to uncover a novel connection between gene expression variation and an organismal trait.

## Materials and Methods

### Strains and growth conditions

Strains and primers used in this study are listed in S2 and S3 Tables, respectively. The parental strains for QTL mapping were YPS163 (oak strain) and the S288c-derived DBY8268 (lab strain; referred to throughout the text as S288c). The construction of the S288c x YPS163 QTL mapping strain panel (44 F_2_ progeny) is described in [96] (kindly provided by Justin Fay). Genotypes for the strain panel are listed in S4 Table. Deletions in the BY4741 (S288c) background were obtained from Open Biosystems (now GE Dharmacon), with the exception of *hap1* (whose construction is described in [45]). Deletions were moved into haploid *MATa* derivatives of DBY8268 (this study) and YPS163 [45] by homologous recombination with the deletion‷KanMX cassette amplified from the appropriate yeast knockout strain [97]. DBY8268 containing a wild-type *HAP1* allele from YPS163 was constructed in two steps. First, the MX cassette from the *hap1*Δ::KanMX deletion was replaced with a URA3MX cassette by selecting for uracil prototrophy. Then, *URA3* was replaced with wild-type *HAP1* from YPS163 (amplified using primers 498-bp upstream and 1572-bp downstream of the *HAP1* ORF), while selecting for loss of *URA3* on 5-fluoroorotic acid (5-FOA) plates. Deletions and repair of *HAP1* were confirmed by diagnostic PCR (see S3 Table for primer sequences). Diploid strains for *HAP1* reciprocal hemizygosity analysis were generated as follows. The hemizygote containing the wild-type S228c *HAP1* allele (JL580) was generated by mating JL140 (YPS163 *MATa ho*Δ::HygMX *hap1*Δ::KanMX) to JL506 (DBY8268 *MATα ho ura3 hap1*). The hemizygote containing the wild-type YPS163 allele (JL581) was generated by mating JL112 (YPS163 *MATα ho*Δ::HygMX *HAP1*) to JL533 (DBY8268 *MATa ho ura3 hap1*Δ::KanMX). All strains were grown in batch culture in YPD (1% yeast extract, 2% peptone, 2% dextrose) at 30°C with orbital shaking (270 rpm).

### Cross protection assays

Cross-protection assays were performed as described in [44] with slight modifications. Briefly, 3-4 freshly streaked isolated colonies (<1 week old) were grown overnight to saturation, sub-cultured into 6 ml fresh media, and then grown for at least 8 generations (>12 h) to mid-exponential phase (OD_600_ of 0.3 – 0.6) to reset any cellular memory of acquired stress resistance [85]. Each culture was split into two cultures and pretreated with YPD media containing either a single mild “primary” dose or the same concentration of water as a mock-pretreatment control. Primary doses consisted of 5% v/v ethanol, 0.4 M NaCl, or 0.4 mM H_2_O_2_. Thereafter, mock and primary-treated cells were handled identically. Following 1-hour pretreatment at 30°C with orbital shaking (270 rpm), cells were collected by mild centrifugation at 1,500 = *g* for 3 min. Pelleted cells were resuspended in fresh medium to an OD_600_ of 0.6, then diluted 3-fold into a microtiter plate containing a panel of severe “secondary” H_2_O_2_ doses ranging from 0.5 – 5.5 mM (0.5 mM increments; 150 μl total volume). Microtiter plates were sealed with air-permeable Rayon films (VWR), and cells were exposed to secondary stress for 2 hours at 30°C with 800 rpm shaking in a VWR^®^ symphony™ Incubating Microplate Shaker. Four μl of a 50-fold dilution was spotted onto YPD agar plates and grown 48 h at 30°C. Viability at each dose was scored using a 4-point semi-quantitative scale to score survival compared to a no-secondary stress (YPD only) control: 100% = 3 pts, 50-90% = 2 pts, 10-50% = 1 pt, or 0% (3 or less colonies) = 0 pts. An overall H_2_O_2_ tolerance score was calculated as the sum of scores over the 11 doses of secondary stress. Raw phenotypes for all acquired stress resistance assays can be found in S5 Table. A fully detailed acquired stress protocol has been deposited to protocols.io under doi dx.doi.org/10.17504/protocols.io.g7sbzne.

### QTL mapping and Heritability Estimates

Phenotyping of the QTL mapping strain panel for basal and acquired H_2_O_2_ resistance was performed in biological duplicate. Because cross-protection assays on the entire strain panel could not all be performed at the same time, we sought to minimize day-to-day variability. We found that minor differences in temperature and shaking speed affected H_2_O_2_ resistance; as a result, we used a digital thermometer and tachometer to ensure standardization across experiments. Moreover, we found that differences in handling time were a critical determinant of experimental variability. To minimize this source of variability, all cell dilutions were performed quickly using multichannel pipettes, and no more than two microtiter plates were assayed during a single experiment. To ensure that replicates on a given day were reproducible, we always included the YPS163 wild-type parent as a reference.

Single mapping scans were performed using Haley-Knott regression [98] implemented through the R/QTL software package [99]. Genotype probabilities were estimated at every cM across the genome using the calc.genoprob function. Significant LOD scores were determined by 10,000 permutations that randomly shuffled phenotype data (i.e. strain labels) relative to the genotype data. The maximum LOD scores for the permuted scans were sorted, and the 99^th^ percentile was used to set the genome-wide FDR at 1%. This resulted in LOD cutoffs of 3.21 for QTL mapping of basal H_2_O_2_ resistance, and 4.05 for acquired H_2_O_2_ resistance.

Broad-sense heritability (*H^2^*) was estimated from the segregant data as described in [71] using a random-effects ANOVA model implemented through the lmer function in the lme4 R package [100]. *H^2^* was estimated using the equation 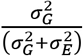 where 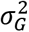 represents the genetic variance due to the effects of segregrant, and 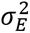 represents the residual (error or environmental) variance.

### Quantitative PCR of *CTT1* expression and cellular peroxidase assays

Induction of *CTT1* by ethanol was assessed by real-time quantitative PCR (qPCR) using the Maxima SYBR q-PCR Master Mix (Thermo Fisher Scientific) and a Bio-Rad CFX96 Touch™ Real-Time PCR Detection System, according to the manufacturers’ instructions. Cells were grown to mid-exponential phase (OD_600_ of 0.3 – 0.6) as described for the cross-protection assays. Cells were collected by centrifugation at 1,500 = *g* for 3 minutes immediately prior to the addition of 5% v/v ethanol (unstressed sample) and 30 minutes post-ethanol treatment, which encompasses the peak of global expression changes to acute ethanol stress [45]. Cell pellets were flash frozen in liquid nitrogen and stored at-80°C until processing. Total RNA was recovered by hot phenol extraction as previously described [101], and then purified with a Quick- RNA™ MiniPrep Plus Kit (Zymo Research) including on-column DNase I treatment. cDNA synthesis was performed as described [101], using 10 μg total RNA, 3 μg anchored oligo-dT (T20VN), and SuperScript III (Thermo Fisher Scientific). One ng cDNA was used as template for qPCR with the following parameters: initial denaturation at 95°C for 3 minutes followed by 40 cycles of 95°C for 15 seconds and 55°C annealing and elongation for 1 minute. Cq was determined using regression analysis, with baseline subtraction via curve fit. The presence of a single amplicon for each reaction was validated by melt curve analysis. The average of two technical replicates were used to determine relative *CTT1* mRNA abundance via the ΔΔCq method [102], by normalizing to an internal control gene (*ERV25*) whose expression is unaffected by ethanol stress and does not vary in expression between S288c and YPS163 [45]. Primers for *CTT1* and *ERV25* were designed to span ~200 bp in the 3’ region of each ORF (to decrease the likelihood of artifacts due to premature termination during cDNA synthesis), and for gene regions free of polymorphisms between S288c and YPS163 (see S3 Table for primer sequences). Three biological replicates were performed and statistical significance was assessed via a paired *t*-test using Prism 7 (GraphPad Software).

For peroxidase activity assays, mid-exponential phase cells were collected immediately prior to and 60 minutes post-ethanol treatment, to assess peroxidase activity levels during the induction of cross protection. Cells were collected by centrifugation at 1,500 = *g* for 3 minutes, washed twice in 50 mM potassium phosphate buffer, pH 7.0 (KP_i_), flash frozen in liquid nitrogen, and then stored at-80°C until processed. For preparation of whole cell extracts, cells were thawed on ice, resuspended in 1 ml KP_i_ buffer, and then transferred to 2- ml screw-cap tubes for bead beating. An equal volume (1 ml) of acid-washed glass beads (425- 600 micron, Sigma-Aldrich) was added to each tube. Cells were lysed by four 30-second cycles of bead beating in a BioSpec Mini-Beadbeater-24 (3,500 oscillations/minute, 2 minutes on ice between cycles). Cellular debris was removed by centrifugation at 21,000 = *g* for 30 minutes at 4°C. The protein concentration of each lysate was measured by Bradford assay (Bio-Rad) using bovine serum albumin (BSA) as a standard [103]. Peroxidase activity in cellular lysates was monitored as described [104], with slight modifications. Briefly, 50 μg of cell free extract was added to 1 ml of 15 mM H_2_O_2_ in KP_i_ buffer. H_2_O_2_ decomposition was monitored continuously for 10 minutes in Quartz cuvettes (Starna Cells, Inc.) at 240 nm (ε_240_ = 43.6 M^−1^ cm^−1^) using a SpectraMax Plus Spectrophotometer (Molecular Devices). One unit of catalase activity catalyzed the decomposition of 1 μmol of H_2_O_2_ per minute. For each sample, results represent the average of technical duplicates. To assess statistical significance, four biological replicates were performed and significance was assessed via a paired *t*-test using Prism 7 (GraphPad Software).

## Acknowledgements

We thank Audrey Gasch and Justin Fay for strains, Andy Alverson and Christian Tipsmark for the use of their equipment, and members of the Lewis lab for helpful conversations. This material is based upon work supported by National Science Foundation Grant No. IOS-1656602 (JAL), startup funds provided by the University of Arkansas (JAL), the Arkansas Biosciences Institute (Arkansas Settlement Proceeds Act of 2000) (JAL), and a Research Assistantship provided through the University of Arkansas Cell and Molecular Biology Graduate Program (ANS). The funders had no role in study design, data collection and analysis, decision to publish, or preparation of the manuscript.

---

**Figure S1.**
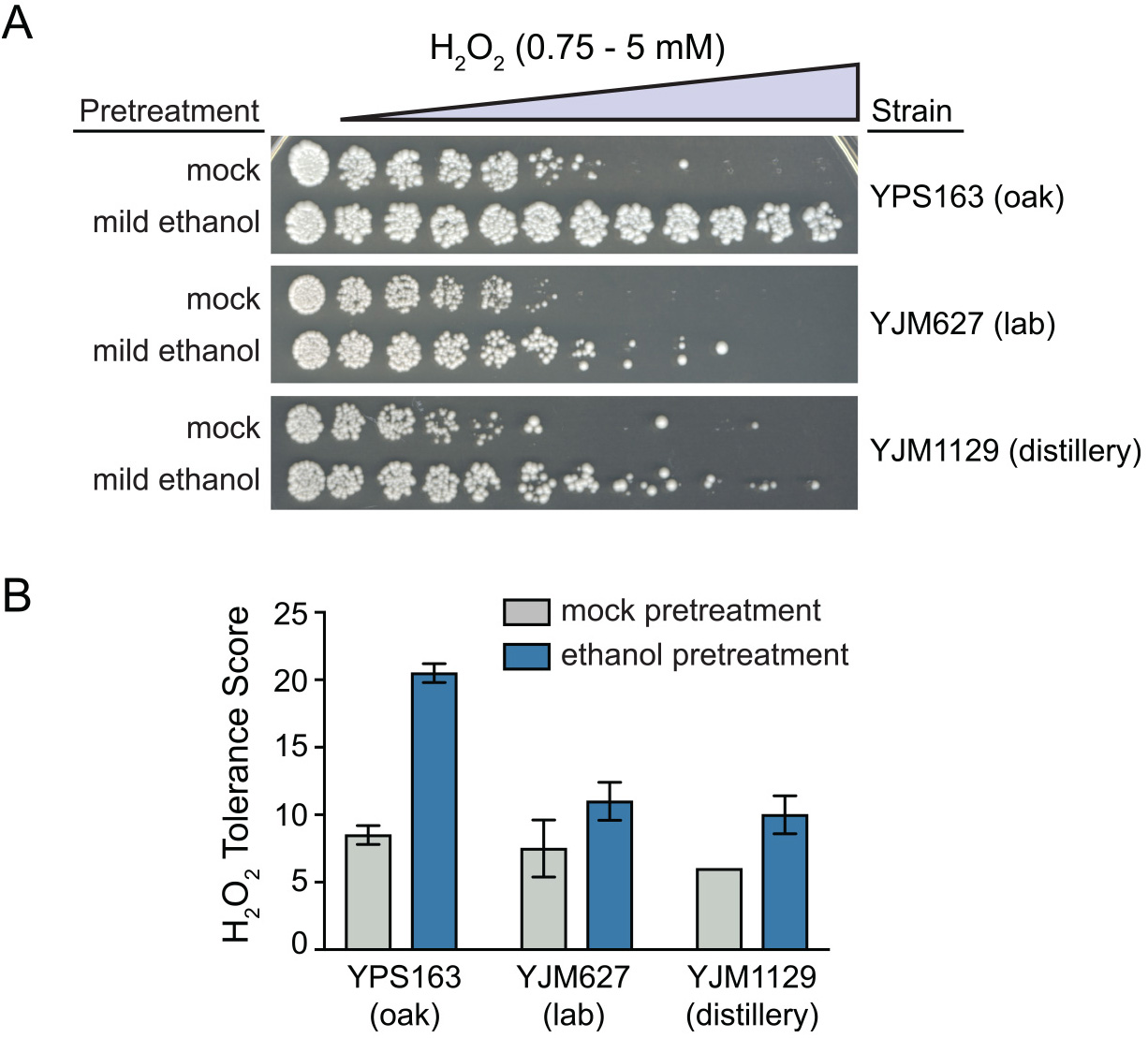
Other non-S288c-derived yeast isolates lack ethanol-induced cross protection against H_2_O_2_. (A) Representative acquired H_2_O_2_ resistance assays for wild-type YPS163, YJM627, and YJM1129. (B) Survival score plots indicating the mean and standard deviation of biological triplicates.

## Supporting information

Table S1. Overlap between genes that are part of the *HAP1* eQTL hotspot, have defective induction by ethanol in S288c vs. YPS163, and are ChIP targets of Hap1p.

Table S2. Strains used in this study.

Table S3. Primers used in this study.

**Table S4. Genotypes for S288c × YPS163 QTL mapping strain panel.** The “Strain” heading for column 1 denotes strain labels for the parental strains (Y = YPS163, S = S288c) and each segregant. Subsequent columns represent genotypes at each marker (Row heading 1 = marker name; Row heading 2 = marker chromosome; Row heading 3 = marker position in cM). Genotypes at each marker are denoted as having the S288c allele (S), YPS163 allele (Y), or missing data (NA).

Table S5. Raw data used to generate each figure.

